# Combined effects of age and BMI are related to altered cortical thickness in adolescence and adulthood

**DOI:** 10.1101/596064

**Authors:** Margaret L. Westwater, Raquel Vilar-López, Hisham Ziauddeen, Antonio Verdejo-García, Paul C. Fletcher

**Author notes:** Please address correspondence regarding this manuscript to: Margaret L. Westwater Douglas House 18b Trumpington Road Cambridge CB1 8AH United Kingdom.

## Abstract

Overweight and obesity are associated with functional and structural alterations in the brain, but how these associations change across critical developmental periods remains unknown. Here, we examined the relationship between age, body mass index (BMI) and cortical thickness (CT) in healthy adolescents (n=70; 14 – 19 y) and adults (n=75; 25 – 45 y). We also examined the relationship between adiposity, impulsivity, measured by delay discounting (DD), and CT of the inferior frontal gyrus (IFG), a region key to impulse control. A significant age-by-BMI interaction was observed in both adolescents and adults; however, the direction of this relationship differed between age groups. In adolescents, increased age-adjusted BMI Z-score attenuated age-related thinning globally and in the right superior frontal gyrus. In adults, increased BMI augmented age-related CT reductions, both globally and in bilateral parietal cortex. Although DD was unrelated to adiposity in both groups, increased DD and BMI were both associated with reduced IFG thickness in adults. Our findings suggest that the known age-related changes in CT in adolescence and adulthood are altered by adiposity. The impact of weight on cortical development and its functional implications would suggest that future studies of adolescent and adult brain development take adiposity into account.

## 1. Introduction

While obesity is defined as excess fat mass (body mass index (BMI) > 29.9 kg/m^2^), it is increasingly recognised as a neurobehavioural condition with metabolic consequences (O’Rahilly and Farooqi, 2008). Body mass is a complex trait with a heritability of 40 – 70% (Maes et al., 1997), and both rare (e.g., *MC4R*; Vaisse et al., 1998; Yeo et al., 1998) and common (e.g. *FTO*) genetic variants relate elevated body mass to altered brain functioning (Locke et al., 2015). In the context of the obesity pandemic, these findings have spurred neuroimaging research into health-harming weight gain. Structurally, elevated BMI has been consistently related to reduced grey matter (GM) and white matter (WM) integrity in the adult and ageing brain (Kurth et al., 2013; Medic et al., 2019, 2016; Pannacciulli et al., 2006; Ronan et al., 2016). However, whether adiposity impacts on the developing brain, particularly during critical periods of accelerated GM and WM changes, remains unknown.

This remains a critical area of study as the proportion of overweight children and adolescents has risen significantly in the past three decades (Lobstein et al., 2015), and the effects of early-life weight gain may be particularly pernicious (Ward et al., 2017). Adolescent overweight relates not only to reduced cardiometabolic health (Reilly and Kelly, 2011) but also to poorer psychological outcomes, including increased incidence of depression and problematic alcohol use (Anderson et al., 2007; Fonseca et al., 2009). These findings may be explained, in part, by the effect of adiposity on cortical morphometric features, such as cortical thickness (CT) or grey matter volume (GMV), which undergo dynamic changes throughout adolescence.

Extensive study of normative brain development (see Casey et al., 2000; Fjell et al., 2009; Giedd et al., 1999; Goddings et al., 2014) has shown that both CT and GMV follow a nonlinear trajectory, peaking at 7 to 10.5 years and 8 years of age, respectively, prior to a rapid decline during adolescence (Mills et al., 2016; Shaw et al., 2008). Changes in these MRI-derived measures are thought to reflect reduced synaptic density and continued WM growth, as part of a putative biological mechanism that underlies the increased cognitive and behavioural capacity of emerging adulthood (Hensch, 2004; Knudsen, 2004). Indeed, CT is increasingly used as a ‘biomarker’ of atypical adolescent development, as thickness alterations have been related to neurocognitive traits like impulsivity and intelligence (Schilling et al., 2013; Shaw et al., 2006), attention-deficit hyperactivity disorder outcomes (Shaw et al., 2006) and the onset of depression (Whittle et al., 2014). Cortical thinning decelerates in early adulthood (Zhou et al., 2015) and accelerates again in the sixth and seventh decade of life, in line with healthy ageing (Kemper, 1994; Salat et al., 2004).

Against this growing understanding of structural brain changes across adolescence and their functional implications, it becomes crucial to examine factors that may influence them. The study of GM alterations in overweight youth has largely been limited to group comparisons of GM volume, with few studies examining CT. In the cortex, increased childhood BMI percentile has been related to GMV reductions in frontal and limbic regions (Alosco et al., 2014). Similar results were reported in obese adolescents, where pathological eating behaviour correlated with reduced orbitofrontal cortex (OFC) volume (Maayan et al., 2011). Overweight adolescents with (Yau et al., 2012) and without (Moreno-López et al., 2012) metabolic syndrome have shown reduced hippocampal volume relative to controls. However, examinations of CT have reported diffuse increases in thickness amongst adolescents with an elevated visceral fat ratio (Saute et al., 2016). Furthermore, a large, longitudinal study reported no impact of body mass on CT in typically developing children (Sharkey et al., 2015).

Studies of GM variation in adult overweight and obesity, while more numerous, have produced similarly inconsistent findings. Elevated BMI (Ho et al., 2010; Raji et al., 2010) and obesity (Gunstad et al., 2009; Yokum et al., 2012) have been associated with reduced total GMV; however, body mass has been consistently unrelated to global mean CT. Local GMV reductions have been observed in the OFC and the anterior cingulate cortex with increasing BMI (Raji et al., 2010), and decreased posterior parietal and temporal volumes have been reported in overweight adults (Gunstad et al., 2009; Walther et al., 2009). Studies examining local CT alterations have primarily shown BMI-related thinning in the left inferior parietal cortex, the inferior temporal and insular cortex (Hassenstab et al., 2012; Veit et al., 2014), and the prefrontal cortex, including ventromedial (Medic et al., 2016), superior frontal and orbitofrontal (Marqués-Iturria et al., 2013) regions. Although the structural correlates of neurocognitive phenotypes in obesity remain poorly characterised, reduced thickness in the superior frontal gyrus has been related to poorer performance on an inhibitory control task and higher BMI (Lavagnino et al., 2016).

Taken together, despite emerging evidence of GM reductions with increasing body mass, the association between adiposity and age-related changes in CT has not been directly examined. We therefore aimed to quantify CT changes associated with increasing age and adiposity in otherwise healthy adolescent and adult cohorts. We predicted that age-related thinning would be increased, both globally and locally, in those with greater adiposity, with local effects found in the frontal lobes of the adolescent group and in bilateral temporal, parietal and ventral frontal lobe regions in adults. In another analysis, we examined the functional correlates of any adiposity-related alterations by assessing whether adiposity and performance on a delay discounting task, taken as a measure of impulsivity, related to CT of the inferior frontal gyrus (IFG), a brain region strongly linked to impulse control (Aron et al., 2014). We hypothesised that 1) increased adiposity would relate to poorer inhibitory control, i.e., steeper delay discounting (DD), in both age groups and 2) a significant BMI by DD interaction would be associated with decreased IFG thickness in adults but not in adolescents, as impaired inhibitory control has been linked to protracted thinning of prefrontal regions in adolescence (Shaw et al., 2011).

## 2. Materials & Methods

### 2.1 Participants

Seventy-nine adolescents (M_age_ ± SD, 16.6 ± 1.5 y; 39 female) and 80 adults (33.3 ± 6.3 y; 45 female) were recruited in Granada, Spain via flyers in the university, local newspapers and radio advertisements in the local community. Adolescent participants were also recruited from surrounding schools. Excess-weight and healthy-weight adolescents were recruited according to the International Obesity Task Force and CDC guidelines on overweight and obesity. As BMI is both sex- and age-dependent in children and adolescents, cut-off scores for healthy weight, overweight and obese adolescents were determined by age-adjusted BMI percentile (Cole and Lobstein, 2012).

Adolescents between 12 and 19 years and adults between 20 and 45 years were eligible for the study. Exclusion criteria for both age groups included left-handedness, current or lifetime psychopathology (e.g., diagnosis of a Diagnostic and Statistical Manual of Mental Disorders, 4^th^ Edition (DSM-IV-TR; American Psychiatric Association, 2000) Axis I disorder, use of psychotropic medication, metabolic syndrome (e.g., diabetes, insulin resistance) and, in adults, morbid obesity (BMI ≥ 40 kg/m^2^). Participants were screened for contraindications to MRI scanning (e.g., ferromagnetic implants, claustrophobia, pregnancy). All participants had normal or corrected-to-normal vision and no history of traumatic brain injury or a neurological disorder.

Participants provided written informed consent prior to participation and received monetary compensation for their time. Adolescent participants signed assent forms, and their parents signed parental consent forms. The study was approved by the Ethics Committee for Human Research at the University of Granada, and all procedures were carried out in accordance with the Declaration of Helsinki.

### 2.2 Procedure

Interested individuals completed a telephone screen with a member of the research team to determine their eligibility for the study. Following the phone screening, eligible volunteers completed the Beck Depression Inventory (Beck et al., 1996) and self-report items adapted from the DSM-*5* (American Psychiatric Association, 2013) criteria for binge-eating disorder. Individuals who reported depressive or binge eating symptoms were excluded. Eligible volunteers underwent MRI scanning as a part of a larger study (described by Navas et al., 2016; Verdejo-Garcia et al., 2015), examining cognition and decision-making in obesity. In brief, participants attended an initial study session, where they provided informed consent or assent and completed neuropsychological tasks and questionnaires, before attending a second scanning session. After the first session, five adolescent participants withdrew from the study, and three adults were excluded because of psychopathology symptoms. Seventy-four adolescents and 78 adults underwent scanning. Following quality assurance procedures (see Supplementary Material), 70 adolescents (35 female, 16.7 ± 1.4 y; 35 male, 16.4 ± 1.5 y) and 75 adult (42 female, 32.8 ± 6.3 y; 33 male, 33.9 ± 6.4 y) participants comprised the final sample, and demographic information by age group is shown in Tables 1 and 2.

**Table 1.** Demographic information by weight class in adolescents.

| Feature | Heathy weight (*n* = 35) |  | Overweight (*n* = 17) |  | Obese (*n* = 18) |  |
| --- | --- | --- | --- | --- | --- | --- |
|  | *M* (SD) | Range | *M* (SD) | Range | *M* (SD) | Range |
| Sex | F = 19, M = 16 |  | F = 9, M = 8 |  | F = 7, M = 11 |  |
| Age (y) | 16.7 (1.4) | 14 – 18 | 16.9 (1.6) | 14 – 19 | 16.2 (1.5) | 14 – 19 |
| Education (y) | 10.7 (1.4) | 8 – 12 | 10.9 (1.6) | 8 – 13 | 10.2 (1.5) | 8 – 13 |
| BMI (kg/m<sup>2</sup>) | 21.2 (2.1) | 17.2 – 25.3 | 26.7 (1.4) | 23.9 – 29.4 | 32.3 (2.4) | 26.4 – 36.3 |
| BMI Percentile | 51 (23) | 3 – 84 | 90 (2.7) | 85 – 94 | 98 (1) | 95 - 99 |
| Age-adjusted BMI Z-Score | 0.0 (0.7) | -2.0 – 1.0 | 1.3 (0.2) | 1.0 – 1.6 | 2.1 (0.2) | 1.7 – 2.5 |
| Body fat (%) | 16.1 (7.8) | 3 – 31.3 | 23.4 (10.2) | 3 – 37.9 | 31.5 (9.4) | 14.2 – 44.5 |
| Binge score | 1.2 (1.1) | 0 – 3 | 0.5 (0.7) | 0 – 2 | 1.2 (1.0) | 0 – 3 |
| BDI | 5.7 (5.0) | 0 – 14 | 6.2 (5.8) | 0 – 17 | 4.8 (5.0) | 0 – 14 |
| MCQ *k* | 0.03 (0.03) | 0 – 0.13 | 0.02 (0.03) | 0 – 0.13 | 0.05 (0.08) | 0 – 0.25 |
| MCQ AUC | 0.52 (0.22) | 0.09 – 0.14 | 0.61 (0.26) | 0.14 – 0.98 | 0.56 (0.30) | 0.09 – 0.92 |

**Notes:** F = female; M = male; BMI = Body mass index; AUC = area under the curve; MCQ = Monetary Choice Questionnaire; BDI = Beck Depression Inventory

**Table 2.** Demographic information by weight class in adults.

| Feature | Heathy weight (*n* = 34) |  | Overweight (*n* = 21) |  | Obese (*n* = 20) |  |
| --- | --- | --- | --- | --- | --- | --- |
|  | *M* (SD) | Range | *M* (SD) | Range | *M* (SD) | Range |
| Sex | F = 20, M = 14 |  | F = 11, M = 10 |  | F = 11, M = 9 |  |
| Age (y) | 32.9 (6.5) | 25 – 45 | 35.0 (6.3) | 25 – 45 | 32.2 (2.6) | 25 – 45 |
| Education (y) | 18.6 (3.7) | 12 – 25 | 17.9 (3.6) | 9 – 25 | 17.0 (4.0) | 10 – 24 |
| BMI (kg/m<sup>2</sup>) | 22.1 (1.7) | 19.0 – 24.8 | 27.6 (1.6) | 25.2 – 29.9 | 33.5 (2.6) | 30 – 38.3 |
| Body fat (%) | 20.4 (6.1) | 12.2 – 34.4 | 29.3 (7.7) | 15.7 – 40.9 | 34.6 (8.8) | 16.7 – 44.4 |
| BDI | 5.8 (5.4) | 0 – 16 | 3.4 (4.4) | 0 – 15 | 3.7 (4.3) | 0 – 16 |
| MCQ *k* | 0.02 (0.02) | 0 – 0.11 | 0.02 (0.03) | 0 – 0.13 | 0.01 (0.01) | 0 – 0.04 |
| MCQ AUC | 0.59 (0.21) | 0.14 – 0.93 | 0.50 (0.20) | 0.14 – 0.93 | 0.69 (0.19) | 0.29 – 1 |

**Notes:** F = female; M = male; BMI = Body mass index; AUC = area under the curve; MCQ = Monetary Choice Questionnaire; BDI = Beck Depression Inventory

### 2.3 Measures

#### *Beck Depression Inventory II* (BDI-II; Beck, Steer, Ball, & Ranieri, 1996)

The BDI-II is a 21-item self-report measure of depression, which has been widely used in both clinical and research settings. Participants responded to items indexing depressive symptoms (e.g., changes in sleeping habits, appetite, and feelings of hopelessness) using a 4-point Likert scale that ranged from 0 to 3. Depressive symptomatology was determined from a total sum score.

#### *Monetary Choice Questionnaire* (MCQ; Kirby, Petry, & Bickel, 1999)

The MCQ is a widely used self-report measure of temporal discounting – the tendency to discount the subjective value of a reward as a function of its delay. The 27-item questionnaire assesses individual preferences for ‘smaller, sooner’ and ‘larger, later’ rewards that vary in both their value and time to be obtained (e.g., *“Would you prefer $7 today, or $30 in 7 days?”*). Overall discounting was determined by an area-under-the-curve (AUC) analysis, as this has been shown to represent a univariate measure of discounting (Myerson et al., 2001). Plots of subjective value vs. delay were generated for each subject, and AUC (standardized between 0 and 1) was calculated by integrating the plotted function (Myerson et al., 2001). Lower AUC values represent greater discounting of future rewards.

Although AUC was used as the dependent measure for our analyses, we also calculated the discounting parameter *k*, which is a hyperbolic free parameter that indexes the rate at which a subject devalues rewards as a function of time. Using nonlinear regression, *k* was fit to the following function:

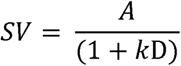

S*V* represents the present value of the reward *A* at delay *D*, and *k* determines the discount rate. As such, increases in *k* correspond with increased discounting of future rewards.

#### Age-adjusted BMI Percentile and Z-Score

For adolescents, BMI percentile and BMI Z-Score for age were calculated based on a standard growth curve validated by the CDC. Values were generated in the EpiInfo 7 toolbox (http://wwwn.cdc.gov/epiinfo/7/), using height (cm), weight (kg), age (days) and sex information.

### 2.3 MRI data acquisition and FreeSurfer reconstruction

Structural MRI data were collected at either 16.00 or 18.00 hours on a Philips 3T scanner, fitted with an eight-channel, phase-array head coil (Intera Achieva, Philips Medical Systems, Eindhoven, The Netherlands). T1-weighted anatomical images were obtained using three-dimensional turbo-gradient-echo sequence (3D-TFE) with sagittal acquisition. This sequence used the following parameters: repetition time (TR) = 8.3 ms, echo time (TE) = 3.8 ms, flip angle = 8°, field of view (FOV) = 240× 240mm, 160 slices with 1 mm^3^ voxels. All MRI scans were reviewed by a neuroradiologist for anatomical abnormalities.

Cortical surface reconstructions were generated in FreeSurfer (v. 5.2; https://surfer.nmr.mgh.harvard.edu/), following an established pipeline (Dale et al., 1999; Fischl et al., 1999). CT was measured at each point, or vertex, on the cerebral surface, representing the shortest mean distance between the WM and pial surfaces. All surface reconstructions were assessed by a blind rater (MLW) who completed minimal white matter and brainmask edits where necessary. Additional information regarding FreeSurfer reconstruction and quality assurance steps can be found in the Supplementary Material.

### 2.4 Global cortical thickness analyses

Multiple linear regression analyses of age, BMI measure, age-by-BMI measure interaction and global CT were completed in R (R Core Team, 2015). We estimated global cortical thickness using the procedure described by Winkler et al. (2010), in the which mean thickness values of each hemisphere are extracted from FreeSurfer, weighted by the corresponding surface area (SA) and summed. By accounting for SA in the calculation of global CT, this approach effectively controls for individual variability in brain size.

Adiposity was modelled continuously in adolescents and adults, using age-adjusted BMI Z-score (henceforth referred to as BMI Z-score) and BMI, respectively. We included sex as a covariate of no interest in all models, and in the adult models, years of education served as a proxy covariate for verbal IQ, which was not assessed in the study. Previous research indicates a strong correlation between educational attainment and verbal IQ in adults (Matarazzo and Herman, 1984), and IQ has been associated with distinct changes in CT throughout adulthood (Narr et al., 2007). As all adolescent participants were enrolled in secondary or tertiary education, years of education was not included as a covariate in these analyses. As such, we tested the following models:

Adolescent models:

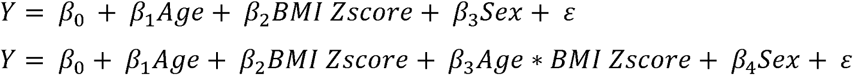

Adult models:

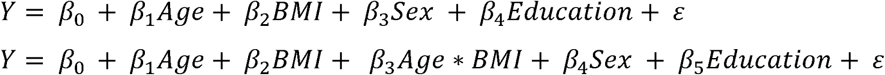

Where Y represents global mean cortical thickness, *β*_*0*_ is the y-intercept, *β*_*1-5*_ are the regression coefficients for each independent variable and ε represents the error term. Because *a priori* hypotheses focused on the interaction of age and weight status specifically, an interaction term was added to the main effect model. For each test, normal distribution of the model residuals was confirmed with the Shapiro-Wilk test.

### 2.5 Vertex-wise cortical thickness analyses

Vertex-wise analyses were performed with the general linear modelling application implemented in FreeSurfer. First, subject surfaces were registered to the fsaverage surface template and smoothed with a 15-mm Gaussian kernel at full width half maximum (FWHM). Next, three GLM matrices were evaluated in each hemisphere of the adolescent and adult groups: 1) correlations with age; 2) correlations BMI measure (BMI Z-score and BMI for adolescents and adults, respectively); and 3) correlations with the interaction of age and BMI measure. Age and BMI measure values were mean-centred before computing the interaction term. As in the global analyses, WM SA and sex were included as covariates of no interest in all vertex-wise analyses; years of education was also entered as a nuisance variable in the adult models. Main and interaction effects were modelled using the “Different Offset, Same Slope (DOSS)” option, which accounts for the possibility of sex differences in the intercepts but assumes fixed slopes of age, BMI measure and the age-by-BMI measure interaction. To correct for multiple comparisons, Monte Carlo Null-Z simulations (10,000 iterations) were applied to the resulting thickness maps to identify clusters of vertex-wise thickness values that significantly differed from zero (alpha level = p < 0.05).

### 2.6 Region of Interest (ROI) analysis of delay discounting, adiposity and IFG thickness

Prior to the ROI analysis, partial correlations between DD (indexed by AUC and the *k* parameters) and BMI measure were assessed in each age group. These correlations controlled for the effects of sex and, in the adult cohort, for years of education (see Section 2.4). Although AUC was selected as the primary measure of discounting for ROI analyses, correlations with the hyperbolic free parameter *k* were also tested, as these can be used as descriptive statistics for comparisons across studies.

Average cortical thickness of the IFG was extracted for each subject using the Destrieux parcellation, which classifies the IFG as the region located between the circular sulcus of the insula and the inferior frontal sulcus that continues into the lateral orbital sulcus (Destrieux et al., 2010). The junction of the inferior portion of the precentral sulcus, the anterior subcentral sulcus and the superior portion of the insular circular sulcus form the posterior boundary. The IFG is further divided into three sections, and, because this region has been functionally implicated in impulse control tasks (Aron et al., 2014; Tabibnia et al., 2011), average thickness of the triangular part was used for ROI analyses.

Sex was included as a covariate in all models, and educational attainment was included in adult models. As we examined IFG thickness in each hemisphere separately, the alpha threshold was reduced to 0.025 (0.05/2) to account for multiple testing.

## 3. Results

### 3.1 Adolescents

#### 3.1.1 Multiple regression analysis of age, BMI Z-Score and global cortical thickness

As expected, there was a significant main effect of age (β = −0.03, t(66) = −4.44, p = 3.46e-05) on global mean CT, where increasing age related to CT. We observed a trend for a main effect of sex (β = −0.04, t(66) = −1.93, p = 0.058) on mean CT, with lower CT in males. The model with age, BMI and sex explained approximately 30% of the variance in global thickness (F(3, 66) = 10.77, p = 7.542e-06, R^2^_*Adjusted*_ = 0.30). A significant age-by-BMI Z-score interaction, where increased body mass related to a greater mean thickness across age (β = 0.01, t(65) = 2.10, p = 0.039), explained an additional 3% of variability in global mean CT (F(4, 65) = 9.59, p = 3.705e-06, R^2^_*Adjusted*_ = 0.33). A nonsignificant three-way interaction confirmed that the age-by-BMI Z-score interaction did not differ between sexes (β = −0.01, t(62) = −0.82, p = 0.42). Thus, increased adiposity was associated with an attenuation of age-dependent cortical thinning across adolescence (Figure 1A).

**Figure 1.**
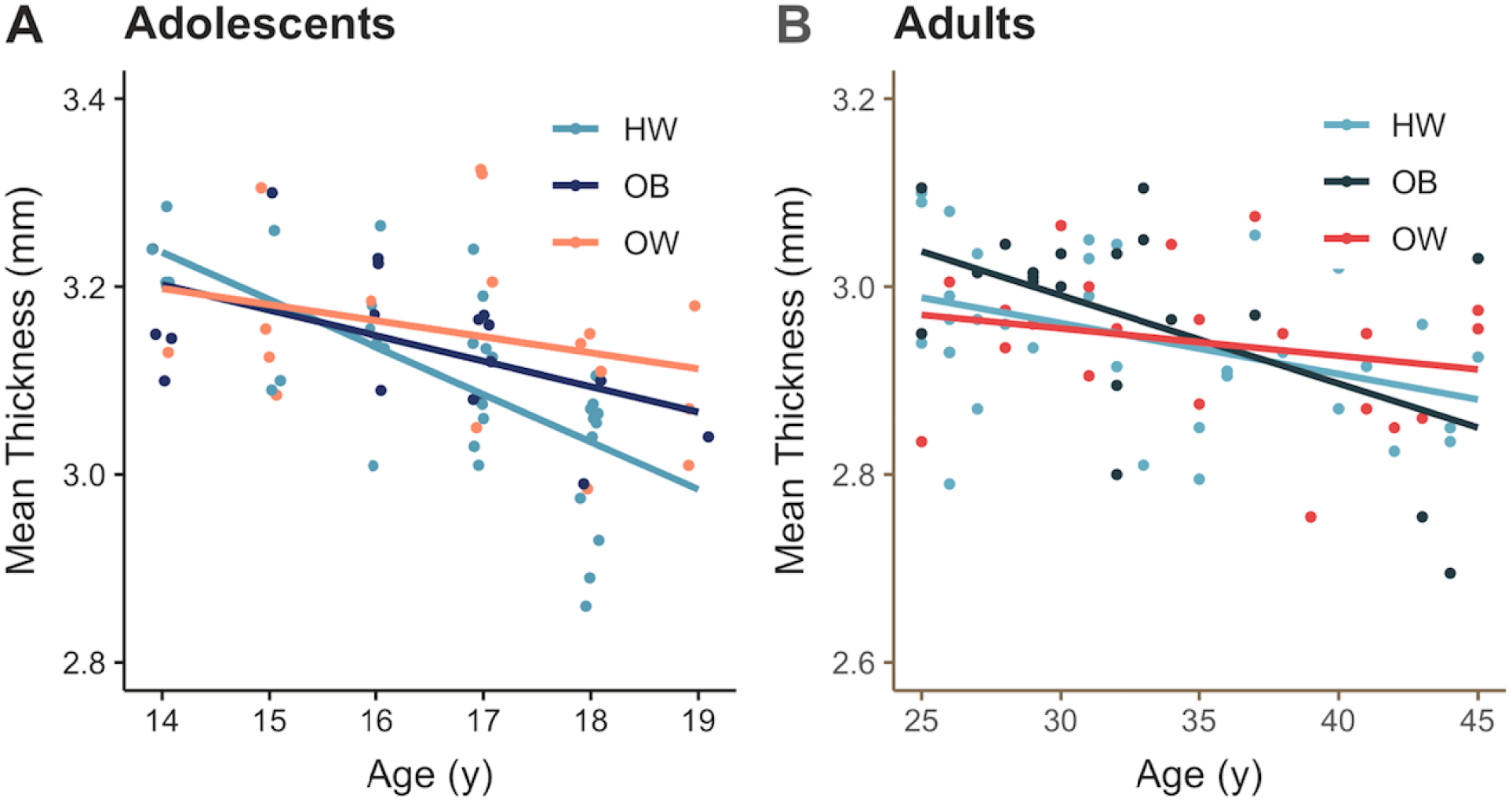
Combined effects of age and BMI-measures on global cortical thickness. **A)** Excess-weight adolescents show increased global CT between ages 14 and 19 relative to lean adolescents. X-axis values were jittered for improved visualisation. **B)** Increasing age and BMI are related to decreased global CT in adults. Data have been plotted by weight group for improved visualisation.

#### 3.1.2 Vertex-wise analysis of cortical thickness

Prior to testing an age-by-BMI Z-score interaction, we generated two-tailed t-statistic maps, examining the main effects of age and BMI Z-score on CT amongst adolescents. Greater BMI Z-scores related to increased CT in the right pericalcarine sulcus and the left lateral occipital cortex (Supplementary Table 2). Moreover, a significant age-by-BMI Z-score interaction showed reduced cortical thinning across age in the right superior frontal gyrus in participants with higher adiposity (Table 3; Figure 2A, C).

**Table 3.** Increased BMI Z-score relates to attenuated superior frontal gyrus cortical thinning across age in adolescents.

| FreeSurfer Region | Side | Cluster Size | | Peak MNI Coordinates | | | Peak Score | $P_{corr}$ |
| --- | --- | --- | --- | --- | --- | --- | --- | --- |
|  |  | Vertices | Size (mm<sup>2</sup>) | X | Y | Z | t |  |
| Superior frontal | R | 7406 | 4341.2 | 19.4 | 33.7 | 43.1 | 2.40 | .0002 |

Notes: p-values < .05 for Monte Carlo Null-Z simulations for multiple comparisons correction.

**Figure 2.**
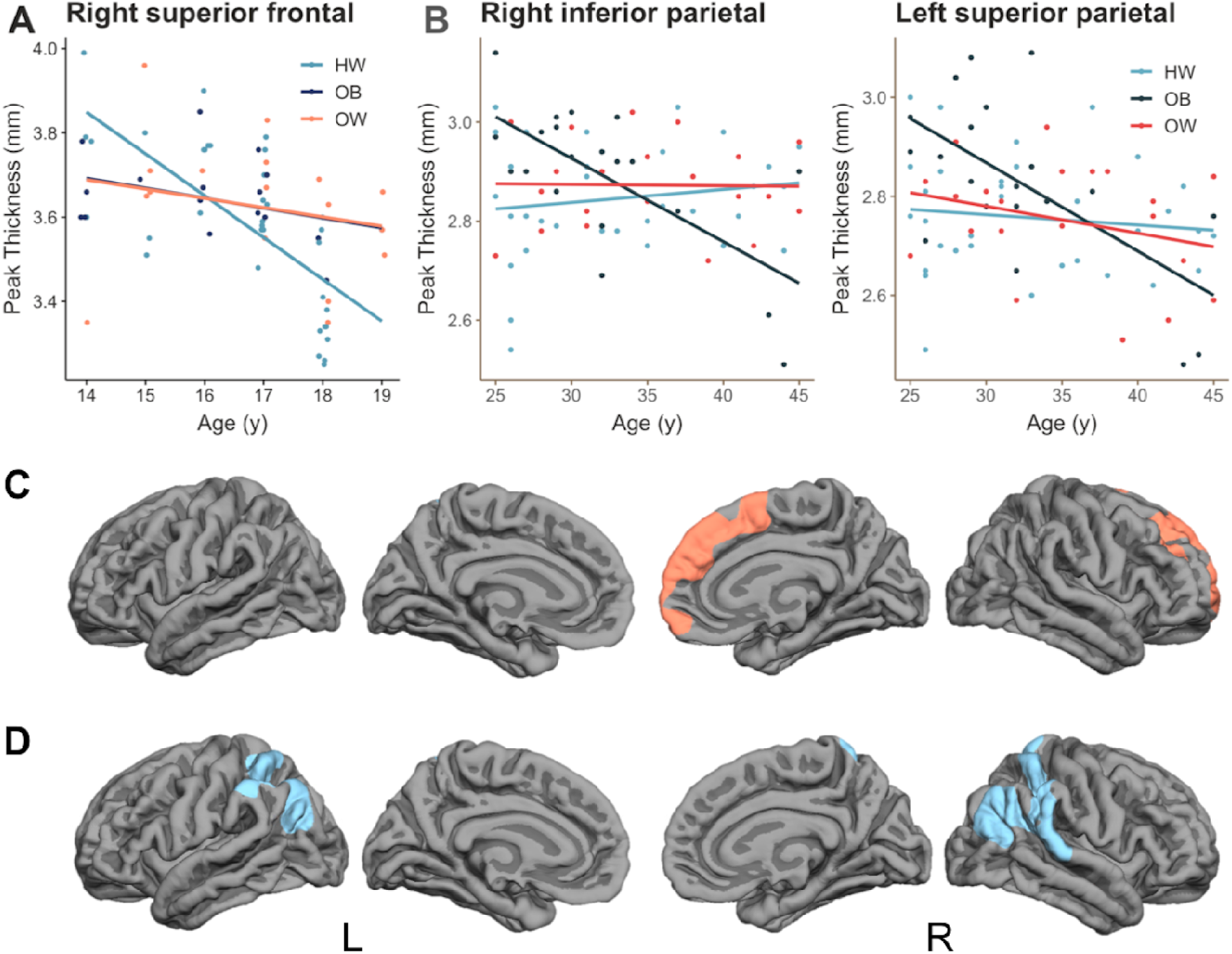
Combined effects of age and adiposity relate to altered local cortical thickness in adolescents and adults. **A & C)** In adolescents, increased BMI Z-score and age related to greater CT in the right superior frontal gyrus. **B & D)** A significant age-by-BMI interaction was associated with reduced bilateral parietal CT in mid-adulthood. All scatterplots show peak CT values of significant clusters (Monte Carlo Null-Z simulation corrected p < .05). To improve visualisation, the continuous age by BMI measure interaction has been plotted by weight group.

#### 3.1.3 ROI analysis of delay discounting, adiposity and IFG thickness

Due to missing data, correlation analyses of BMI Z-score, AUC and *k* parameter included 69 and 67 participants, respectively. BMI Z-Score was not significantly correlated with AUC (p = .56) or *k* parameter (p = .36). Omnibus models examining the relationship between BMI Z-Score, AUC and their interaction were nonsignificant for both left and right IFG CT (all p-values > .20).

### 3.2 Adults

#### 3.2.1 Multiple regression analysis of age, BMI and global cortical thickness

In the adult group, there was a main effect of age (β = −0.007, t(69) = −4.34, p = 4.72e-05) on global mean CT, but the main effect of BMI was nonsignificant (β = 6.00e-04, t(69) = 0.32, p = 0.75). Of the two covariates, only sex exerted a significant effect (β = −0.06, t(69) = - 2.95, p = 0.004) on global thickness. The linear model of age and BMI explained approximately 24% of the variance in global thickness amongst adults (F(4, 69) = 6.85, p = 1.06e-04, R^2^_*Adjusted*_= 0.24). A significant age by BMI interaction (β = −5.94e-04, t(68) = −2.14, p = 0.036) explained an additional 4% of the variance in global mean CT (F(5, 68) = 6.67, p = 4.12e-05, R^2^_*Adjusted*_ = 0.28). For completeness, a three-way interaction between age, BMI and sex was tested, and the age by BMI interaction did not differ across sexes (β = 5.23e^-4^, t(66) = 0.97, p = 0.34). As with the adolescent cohort, adiposity was associated with a significant alteration in age-related cortical thinning (Figure 1B); however, in adults, this effect was in the opposite direction, where adiposity related to greater reductions in CT.

#### 3.2.2 Vertex-wise analysis of cortical thickness

Vertex-wise thickness maps indicated widespread cortical thinning in the frontal, temporal and parietal lobes that was associated with increasing age (Supplementary Table 3). The main effect of BMI on local CT was nonsignificant. However, the interaction of age and BMI was significantly related to CT in the left superior parietal lobule and right inferior parietal lobule, extending to the superior temporal gyrus (see Table 4, Figure 2B, D), where adiposity augmented age-related CT reductions.

**Table 4.** Increased age and BMI relates to reduced cortical thickness in adults.

| FreeSurfer Region | Side | Cluster Size | | Peak MNI Coordinates | | | Peak Score | $P_{corr}$ |
| --- | --- | --- | --- | --- | --- | --- | --- | --- |
|  |  | Vertices | Size (mm<sup>2</sup>) | X | Y | Z | t |  |
| Inferior parietal | R | 13741 | 6557.8 | 41.8 | -72.9 | 22.7 | -5.95 | .0002 |
| Superior parietal | L | 7446 | 3434.8 | -26.9 | -49.3 | 61.0 | -3.42 | .001 |

Notes: p-values < .05 for Monte Carlo Null-Z simulations for multiple comparisons correction.

#### 3.2.3 ROI analysis of delay discounting, adiposity and IFG thickness

Two adults failed to complete the MCQ, and *k* parameter estimates were unavailable for three additional volunteers, leaving 73 and 71 volunteers for behavioural analyses of AUC and *k* parameter, respectively. Neither AUC (p = .46) nor *k* parameter (p = .83) were correlated with BMI.

Two participants who were identified as outliers (CT values 3 SD above or below the mean) were excluded from left IFG analyses; the final sample included 71 volunteers. The main effects of BMI and DD on CT in the left and right IFG were nonsignificant. Age was negatively related to both left (β = −0.01, t(65) = −3.39, p = .001) and right (β = −0.02, t(67) = −4.30, p = 5.70e-05) IFG thickness. Sex was significantly related to right IFG thickness (β = −0.12, t(67) = −2.39, p = .02). A significant BMI by DD interaction indicated reduced IFG thickness with increasing BMI and increasing discounting (decreasing AUC); however, this effect was limited to the left hemisphere (β = −0.06, t(64) = −3.27, p = .002; Figure 3). The main effect model accounted for approximately 20% of the variability in left IFG thickness (F(5, 65) = 4.40, p = .002, R^2^_*Adjusted*_ = 0.20), and the interaction captured an additional 10% of the variance in CT (F(6, 64) = 5.99, p = 5.18e-05, R^2^_*Adjusted*_ = 0.30).

**Figure 3.**
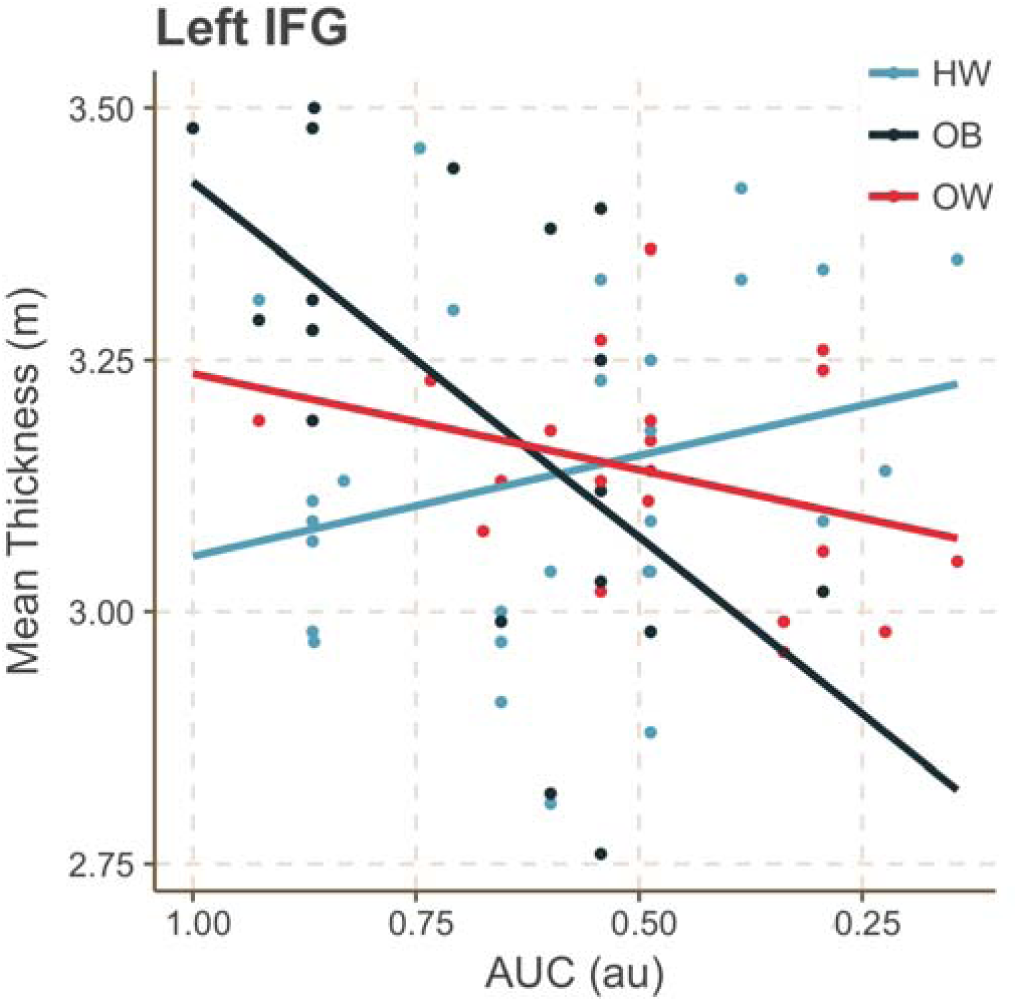
Delay discounting is related to decreased IFG thickness in high-BMI adults. DD moderated the relationship between BMI and left IFG thickness; adults with elevated BMI and DD had reduced IFG thickness. Area under the curve (AUC) was used to measure degree of discounting, where lower AUC represents higher DD. For illustration purposes, data have been plotted by weight group (HW = healthy weight, OB = obese, OW = overweight).

## 4. Discussion

Given increasing evidence for the neurobehavioural causes and consequences of obesity, it will be important to establish, as precisely as possible, the brain alterations associated with excess weight. However, examination of such alterations is necessarily superimposed on the non-linear developmental trajectory of the brain, and therefore, a comprehensive understanding of weight-related alterations in the cortex must consider interactions with these underlying growth processes. The current study provides a potentially important insight into how weight status relates to adolescent and adult brain development in different ways. Using structural MR images from adolescents and adults with no known clinical comorbidity, we found, both in global and local analyses, that the expected cortical thinning is attenuated by increasing adiposity in adolescents but augmented by elevated BMI in adults.

The delay discounting task provides some limited information about the functional relevance of the observed interactions between age and body mass. In adults, but not adolescents, increasing BMI and temporal discounting of monetary reward (i.e. a greater valuation of smaller immediate rewards over larger delayed reward) related to a reduction in left IFG thickness. This suggests that greater impulsive tendencies in higher BMI adults relates to a structural change in prefrontal cortex.

In line with known trajectories of cortical development, we found cortical thinning with increasing age across adolescence. The attenuation of this effect with increasing BMI Z-score could suggest that increased adiposity modulates the progression of MRI-derived cortical thinning in this critical period. A previous prospective study of typically developing children reported a nonsignificant association between BMI Z-score and CT (Sharkey et al., 2015), leading the authors to postulate that the cortical thinning previously observed with elevated BMI only occurs after adolescence. Indeed, our findings support the hypothesis of developmentally-bound brain alterations in obesity. Moreover, our complementary finding from the local, vertex-wise analysis, suggests a notably strong effect in the right superior frontal gyrus (SFG), extending into the PFC. Both histological and *in vivo* evidence indicate protracted development of anterior regions of the prefrontal cortex (PFC; Brodmann area 10), with CT diminishing at a relatively slower rate than posterior regions (for a review, see Dumontheil et al., 2008). Delayed PFC development is thought to contribute to risky decision-making, poor inhibitory control and difficulties with multitasking in adolescence (reviewed in Blakemore & Robbins, 2012), and it has been reported in clinical conditions marked by impulsivity, namely ADHD (Shaw et al., 2007). Moreover, fMRI research of inhibitory control has shown that mean PFC signal intensity increases from adolescence into emerging adulthood (Rubia et al., 2000; Rubia et al., 2006). Given the diverse functional role of the SFG in executive function, cognitive control and sensorimotor processes (Li et al., 2013), delayed cortical maturation of the region may affect cognitive control of food intake, leaving excess-weight adolescents vulnerable to overconsumption despite negative health outcomes (e.g., cardiometabolic conditions).

In contrast to the adolescent group, the negative main effect of age on global and local thickness estimate was exacerbated in adults with increased BMI. We did not observe a significant main effect of BMI on CT, which may relate to the exclusion of individuals with class III obesity (BMI > 39.9) and the subsequently restricted BMI range of the sample. Alternatively, the absence of a main effect of BMI aligns with the observed age-by-BMI measure interaction in both age cohorts: as a consequence of attenuated cortical thinning in adolescence, excess-weight adults have increased cortical thickness in early adulthood that declines significantly with age. Such a progression would obfuscate any main effect of BMI. Nonetheless, our findings accord with previous reports of accelerated brain atrophy with increasing fat mass in adulthood (e.g., Ronan et al., 2016; Walhovd et al., 2014). We found that increased adiposity across age related to reductions in bilateral parietal thickness, and thinning of these regions has been associated with normative ageing in the fifth and sixth decades of life (Fjell et al., 2009).

Moreover, Vainik et al. (2018) reported a positive association between BMI and temporo-parietal thickness, primarily in the right hemisphere, in young adults (M_age_ = 28.85). Interestingly, in our adolescent cohort, BMI Z-Score related to increased thickness of the right pericalcarine sulcus and the left lateral occipital cortex. Medic and colleagues (2016) further reported thinning of the lateral occipital cortex with increasing BMI in middle adulthood. These findings broadly align with theories of altered visuospatial processing in obesity, where external cues may override interoceptive signals (Schachter, 1968); however, the relationship between parietal thickness, age effects and visuospatial ability has not been tested explicitly in obese groups.

We examined associations between delay discounting, and thickness of the IFG, a cortical area thought to enable braking or full stopping of response tendencies (Aron et al., 2014). Contrary to previous reports of increased delay discounting in obesity (e.g., Rasmussen et al., 2010), neither BMI Z-Score nor BMI were significantly correlated with discounting of monetary reward in adolescents and adults. However, amongst adults, the interaction of BMI and delay discounting (i.e., AUC) was related to reduced thickness of the left IFG. Although the right IFG has primarily been implicated in response inhibition, cognitive testing of patients with well-defined left IFG lesions indicates that structural integrity of both nodes is critical for successful motor suppression (Swick et al., 2008). BMI has previously been associated with inferior frontal thinning (Vainik et al., 2018), and in a separate study, reduced functional activity in the inferior frontal cortex predicted weight gain amongst overweight women who demonstrated steeper discounting of monetary reward (Kishinevsky et al., 2012). In addition, Lavagnino and colleagues (2016) reported that, in obese adults, SFG thickness was negatively related to both BMI and reaction time on an inhibitory control task, and reaction time fully mediated the CT-BMI relationship. However, given the dearth of longitudinal studies on cognitive control, obesity and cortical morphometry, causal inferences should be made cautiously. One possible interpretation aligns with the last-in, first-out hypothesis of cortical development (Raz, 2000): the proposed delay of prefrontal development in adolescent overweight corresponds with early atrophy in mid-adulthood overweight, particularly amongst individuals with poorer impulse control.

We can only speculate, of course, on the mechanisms linking altered profiles of cortical thickness changes with increasing BMI. It may be that the observed CT alterations relate to peripheral metabolic features of diet-induced weight gain, specifically increased white adipose tissue. Increases in adiposity result in elevations in leptin, a satiety hormone with central effects on hypothalamic neurons (Considine et al., 1996). Leptin also acts upon peripheral targets that modulate various aspects of physiological functioning, including skeletal growth, pubertal onset and neural development (Ahima et al., 2000; Nagatani et al., 1998). However, while the overproduction of leptin initially serves as a negative feedback signal to reduce energy intake, sustained elevation of the adipokine ultimately leads to leptin resistance, emerging from defective leptin transport across the blood brain barrier and changes to leptin-responsive neurons (El-Haschimi et al., 2000; Shek and Scarpace, 2000). Currently, animal model studies provide the strongest evidence of neurotrophic properties of leptin, reporting increased neurogenesis (Ahima et al., 2011; Udagawa et al., 2006) and glial cell proliferation (Udagawa et al., 2007) with elevated leptin levels in early mammalian development. In humans, leptin replacement therapy has led to sustained increases in GMV of the anterior cingulate cortex, inferior parietal lobe and cerebellum (Matochik et al., 2009). Although the extent to which leptin alters GM architecture cannot be directly determined using voxel-based morphometry, as altered GMV could reflect neuronal cell volume, altered focal myelination, dendritic spine growth or a combination of the three, these findings provide critical evidence of leptin-dependent changes in the cortex. Leptin levels have been shown to increase with age, and some have argued that ageing, along with obesity, constitutes a state of leptin resistance (Lee, 2011). While very preliminary, our findings might suggest that the neurotrophic and neuroprotective effects of leptin are modulated in overweight and obesity. We encourage future research to directly measure leptin levels to determine their effect on the excess weight—cortical thickness relationship across development.

Despite several notable strengths, our findings should be interpreted in light of several limitations. First, we examined cross-sectional, structural MRI data, and prospective studies will be critical to disentangling associations between obesity and cortical thickness across the lifespan. These designs would enable the use of alternative analytic techniques, such as growth mixture modelling, to examine nonlinear maturation of CT in overweight and obesity. A second limitation is the modest sample size. Adiposity likely yields a small effect on cortical thickness, and according to a recent examination of statistical power in cross-sectional structural MRI (Pardoe et al., 2013), larger samples will be necessary to detect thickness changes less than 0.25 millimetres. Third, due to the age- and sex-dependent nature of body mass during childhood, we were unable to directly compare adiposity and age effects in the adolescent and adult cohorts, and future investigation should leverage more precise estimates of fat mass (e.g., those derived from dual-energy X-ray absorptiometry scanning) that can be acquired across the lifespan. Fourth, pubertal onset, lifetime weight trajectories and the interaction of these were not available in this study, but such data would help address the question of whether obesity affects cortical structure differently at certain points in development. Finally, we measured impulse control using monetary delay discounting, and future work would benefit from the use of impulsivity tasks with clinically-relevant rewards (e.g., food) to demonstrate convergent validity with the MCQ.

## 5. Conclusion

Our findings demonstrate that adiposity is associated with significant alterations in the trajectories of cortical change across adolescence and adulthood. Strikingly, these alterations were in opposite directions across the two periods. We observed increased thickness of prefrontal cortical regions for age amongst overweight and obese adolescents, which may leave these individuals vulnerable to maladaptive energy intake as they transition into adulthood. Moreover, we found that the combined effects of elevated BMI and steeper delay discounting are related to lower left IFG thickness, further supporting the view of obesity as a neurobehavioural syndrome. These findings underscore the complex relationship between adiposity and brain health, stressing that any impact of weight on brain structure must be considered in terms of underlying trajectories of age-related change. Furthermore, they strongly suggest that assessments of adolescent developmental trajectories take adiposity into account.

## Acknowledgements

This work was supported by the Bernard Wolfe Health Neuroscience Fund (HZ, PCF) and the Andalusian Health Service (Consejeria de Salud) [project grant P-10-HUM-6635 (NEUROECOBE)] (AVG). MLW was supported by the Cambridge Trust and NIH-Oxford Cambridge Scholars’ Program.

