## Supplementary material for "Combined effects of age and BMI are related to altered cortical thickness in adolescence and adulthood"

**Methods**

*Participants.* Lean adolescents were classified as those with 5^th^ ≤ age-adjusted BMI < 85^th^ percentile; overweight adolescents were those with 85^th^ ≤ age-adjusted BMI < 95^th^ percentile; and obese adolescents had adjusted BMI scores ≥ 95^th^ percentile. In the adult group, lean individuals were identified by 18.5 ≤ BMI ≤ 24.9 kg/m^2^; overweight adults 25.0 ≤ BMI ≤ 29.9 kg/m^2^; and obese adults ≤ 30.0 BMI < 40.0 kg/m^2^, the threshold for morbid obesity. To confirm the absence of metabolic comorbidities, blood samples were acquired from a subset (n = 43) of adolescents and assayed for insulin and total cholesterol levels. Samples were collected between 8.00 and 10.00 following an overnight fast.

*Structural MRI Analyses.* Structural MRI data were analysed using FreeSurfer (v 5.2, <https://surfer.nmr.mgh.harvard.edu/>) software. Briefly, the FreeSurfer pipeline first corrects MR images for magnetic field inhomogeneities. Normalized intensity images are created, affine-registered to the Talairach atlas (Talairach & Tournoux, 1988) and skull-stripped to remove non-brain tissue. In the preliminary segmentation, WM voxels are identified by their location, their intensity and the intensity of their adjacent voxels. The two hemispheres are separated, and WM voxels are grouped using a six-neighbours connected components algorithm, thus creating a single-filled volume for each hemisphere. A mesh of triangular faces is fitted to the resulting surface, with two triangles for each exposed voxel face, and it is then smoothed to reduce distortions. The initial smoothing procedure employs an algorithm that accounts for the local intensity of the original images (Dale & Sereno, 1993) at a higher spatial resolution, using trilinear interpolation. Correction for topological defects described by Fischl et al. (2001) ensures that the surface maintains the same topological properties as a sphere.

Following this initial reconstruction of the surface, a second iteration of smoothing is applied, which results in a realistic representation of the GM/WM boundary. This surface, called the white surface, is then deformed outwards toward the point where tissue contrast is maximal, maintaining the original constraints on smoothness and possibility of self-intersection (Fischl & Dale, 2000). This surface is referred to as the pial surface, as it represents the grey matter boundary that sits below the pia mater. Complete cortical surface reconstructions were visually inspected for quality control purposes and discarded when necessary. (see Table 1).

Surfaces were then parcellated into distinct cortical regions using an automated procedure (Fischl et al., 2004). Within each subject, the cortical surface was transformed to a sphere in order to establish a coordinate system on the ‘unfolded’ surface (Fischl, Sereno, & Dale, 1999). Each vertex was assigned an anatomical label based on: 1) the probability that a given vertex belongs to a certain label, using a Bayesian approach; 2) mean curvature information of the region; and 3) contextual information, which, for each vertex, considers the labels that were assigned to neighbouring vertices. This labelling process iterates until the label assignment of each vertex no longer changes. We used the parcellation scheme developed by Desikan et al. (2006), which includes labels for gyri, sulci and 66 regions of interest. Following this, cortical thickness values for each ROI were calculated as the average distance between the white and pial surface within the region. Calculation of surface area was completed in the subjects’ native space, and values were defined as the sum of the areas of each triangular face located within the ROI. Global CT and SA values in each hemisphere were calculated as averages and summed totals of the ROIs, respectively.

*Quality Assurance.* Surface reconstructions were visually assessed by a blind rater (MLW), classified according to a 5-point Likert scale (0 = poor, 5 = good) and discarded when necessary. Within the adolescent group, 2 (2.7%) reconstructions were of poor quality, 1 (1.4%) was adequate poor, 7 (9.6%) were adequate, 19 (26%) were adequate good and 44 (60.3%) were of good quality. Surface reconstructions within the adult group were rated: 2 (2.6%) poor, 1 (1.3%) adequate poor, 10 (12.8%) adequate, 35 (44.9%) adequate good and 30 (38.5%) good. We discarded three, poor-quality MRI scans from the adolescent group, and a fourth adolescent was excluded because of a corrupt file. Three adult structural scans were discarded for quality concerns. In total, 70 adolescents and 75 adults with high-quality structural scans were included in the CT analyses.

Bland and Altmann plots (Supplementary Figure 1) were used to assess potential measurement bias from quality assurance procedures. The mean difference in cortical thickness approximated zero for adolescent left (M, SD = -0.01, .02 mm) and right (0, .02 mm) hemispheres, as well as bilateral adult cortical thickness (left = -.01, .01 mm; right = -.01, .02 mm).

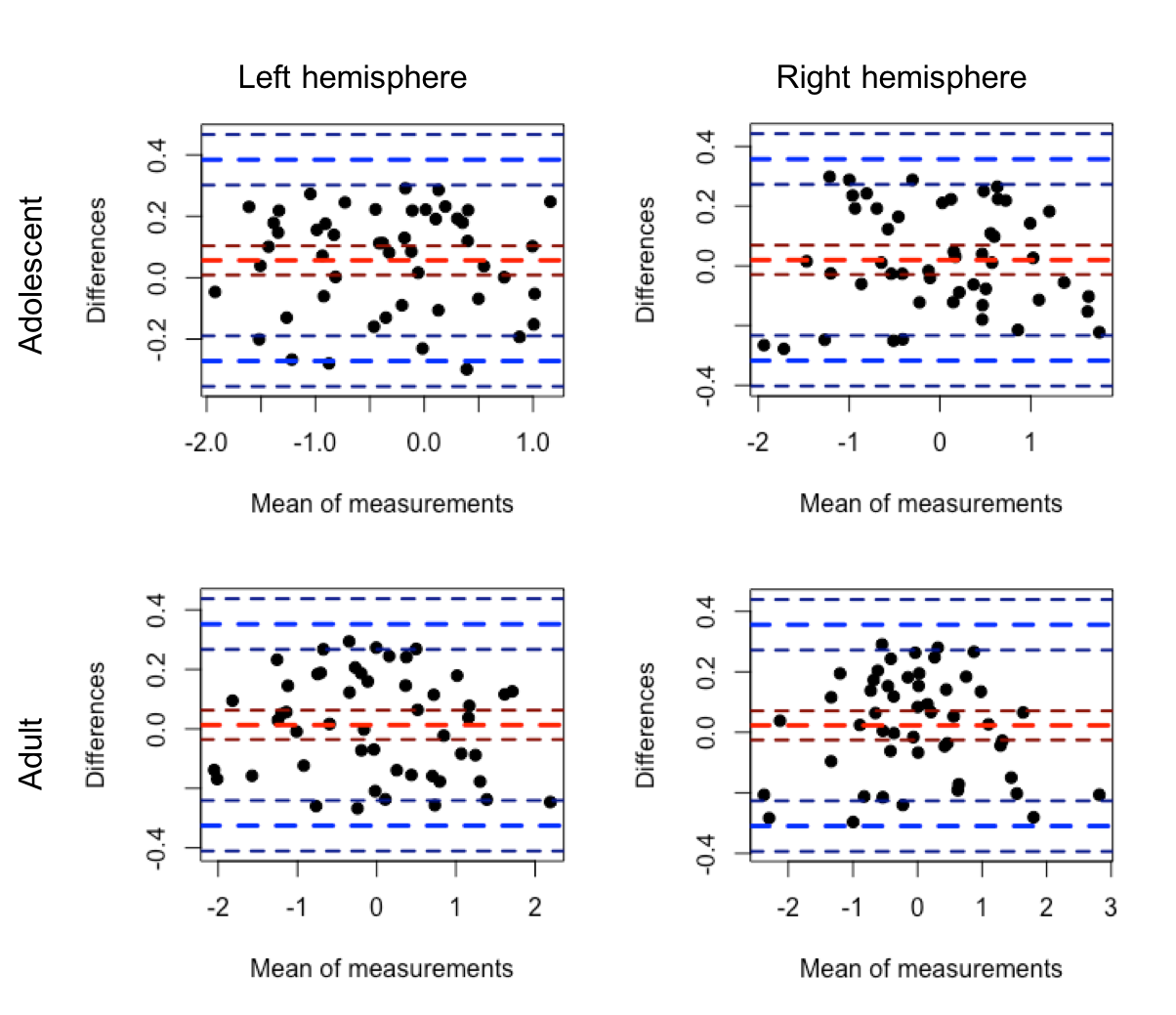

**Supplementary Figure 1.** Bland-Altman plots of pre- and post-edit global cortical thickness values in adolescent and adult groups. Results are presented by hemisphere. The x-axis represents the mean of pre- and post-edit cortical thickness values, and the y-axis illustrates the difference between these estimates.

**Results**

**Supplementary Table 1.** *Correlation between age and cortical thickness in adolescents*

| **#** | **FreeSurfer Region** | **Side** | **Cluster Size** | | **Peak MNI Coordinates** | | | **Peak Score** | **P*_corr_*** |
| --- | --- | --- | --- | --- | --- | --- | --- | --- | --- |
|  |  |  | **Vertices** | **Size (mm^2^)** | **X** | **Y** | **Z** | **T** |  |
| 1 | Precentral | R | 31,991 | 15,714.34 | 17.5 | -9.8 | 60.3 | -8.00 | .017 |
| 2 | Supramarginal | L | 9409 | 4548.32 | -53.4 | -53.3 | 22.6 | -7.74 | .0002 |
| 3 | Pars orbitalis | L | 2223 | 1288.83 | -43.4 | 43.3 | 11.8 | -4.41 | .002 |
| 4 | Lateral orbitofrontal | R | 3759 | 2407.96 | 24.9 | 36.6 | -11.5 | -4.13 | .0002 |
| 5 | Inferior parietal | R | 2996 | 1325.38 | 47.6 | -46.4 | 19.6 | -4.15 | .002 |
| 6 | Cuneus | R | 2578 | 1441.70 | 9.0 | -81.7 | 33.4 | -4.03 | .001 |
| 7 | Superior temporal | R | 2285 | 993.06 | 66.6 | -24.6 | 2.3 | -3.99 | .017 |
| 8 | Postcentral | L | 11,981 | 5189.06 | -44.5 | -19.3 | 16.0 | -3.79 | .0002 |
| 9 | Precentral | R | 2732 | 1605.81 | 19.5 | -50.1 | -8.3 | -3.76 | .0006 |
| 10 | Pars opercularis | L | 1846 | 936.44 | -44.0 | 16.9 | 20.8 | -3.75 | .02 |
| 11 | Cuneus | L | 1629 | 858.24 | -18.7 | -67.2 | 11.3 | -2.66 | .03 |

Notes: p-values < .01 for Monte Carlo Null-Z simulations for multiple comparisons correction.

**Supplementary Table 2.** *Increased BMI Z-score relates to increased cortical thickness in occipital lobe*

| **FreeSurfer Region** | **Side** | **Cluster Size** | | **Peak MNI Coordinates** | | | **Peak Score** | **P*_corr_*** |
| --- | --- | --- | --- | --- | --- | --- | --- | --- |
|  |  | **Vertices** | **Size (mm^2^)** | **X** | **Y** | **Z** | **t** |  |
| Pericalcarine | R | 1566 | 1233.1 | 15.3 | -92.1 | -3.4 | 4.27 | .004* |
| Lateral occipital | L | 7118 | 4814.9 | -22.6 | -98.2 | 1.4 | 3.02 | .0002 |

Notes: p-values < .05, *p-values < .01 for Monte Carlo Null-Z simulations for multiple comparisons correction.

**Supplementary Table 3.** *Correlation between age and cortical thickness in adults*

| **#** | **FreeSurfer**  **Region** | **Side** | **Cluster Size** | | **Peak MNI Coordinates** | | | | **Peak Score** |
| --- | --- | --- | --- | --- | --- | --- | --- | --- | --- |
|  |  |  | **Vertices** | **Size (mm^2^)** | **X** | **Y** | **Z** | **t** | |
| 1 | Caudal middle frontal | R | 13,558 | 7033.0 | 36.7 | 8.0 | 56.0 | -4.61 | |
| 2 | Caudal middle frontal | L | 5010 | 2583.4 | -34.4 | 16.0 | 25.5 | -4.28 | |
| 3 | Cuneus | R | 1367 | 1034.8 | 4.1 | -72.8 | 19.8 | -3.94 | |
| 4 | Inferior parietal | L | 2035 | 1204.9 | -34.9 | -79.2 | 33.1 | -3.73 | |
| 5 | Precentral | L | 3101 | 1321.9 | -32.7 | -22.0 | 43.2 | -3.68 | |
| 6 | Middle temporal | R | 2353 | 1309.7 | 50.4 | 6.2 | -34.6 | -3.54 | |
| 7 | Superior frontal | R | 5000 | 3051.3 | 12.5 | 34.7 | 27.4 | -3.16 | |
| 8 | Middle temporal | L | 2658 | 1565.0 | -49.8 | -60.2 | 10.0 | -3.07 | |
| 9 | Superior frontal | L | 5057 | 3097.4 | -11.3 | 46.1 | 39.4 | -2.96 | |
| 10 | Superior parietal | L | 1763 | 799.1 | -37.4 | -47.1 | 55.0 | -2.52 | |

Notes: p-values < .01 for Monte Carlo Null-Z simulations for multiple comparisons correction.
